# The functional properties of synapses made by regenerated axons across spinal cord lesion sites

**DOI:** 10.1101/2021.06.21.449247

**Authors:** David Parker

## Abstract

While the anatomical properties of regenerated axons across spinal cord lesion sites have been studied extensively, little is known of how the functional properties of regenerated synapses compare to those in unlesioned animals. This comparison has been performed here in the lamprey, a model system for spinal injury research, in which functional locomotor recovery after spinal cord lesions is associated with axonal regeneration across the lesion site.

Regenerated synapses below the lesion site did not differ to synapses from unlesioned axons with respect to the amplitude and duration of single excitatory postsynaptic potentials (EPSPs). They also showed the same activity-dependent depression over spike trains. However, regenerated synapses did differ to unlesioned synapses as the estimated number of synaptic vesicles was greater and there was evidence for an increased postsynaptic quantal amplitude. For axons above the lesion site, the amplitude and duration of single synaptic inputs also did not differ significantly to unlesioned animals. However, in this case there was evidence of a reduction in release probability and inputs facilitated rather that depressed over spike trains.

Synaptic inputs from single regenerated axons below the lesion site thus do not increase in amplitude to compensate for the reduced number of descending axons after functional recovery. However, the postsynaptic input is maintained at the unlesioned level using different synaptic properties. Conversely, the facilitation from the same initial amplitude above the lesion site will make the synaptic input over spike trains functionally stronger. This may help to increase propriospinal activity across the lesion site to compensate for the lesion-induced reduction in supraspinal inputs.

## Introduction

Regeneration remains the dominant approach in attempts to restore function after spinal cord injury (Ramer et al 2014). The anatomical properties of regenerated axons (their number and projections) have been studied extensively in various systems, but their physiological properties have received relatively little attention. Given that these properties will determine the functional effect of any regenerated inputs it is important that they are considered, not least because this may help to explain the variable relationship between regeneration and recovery (Steward et al 2012).

The small size of regenerated axons makes them difficult to study in the mammalian spinal cord. This study has used the lamprey, a model system for studying axonal regeneration and functional recovery after spinal cord lesions. Analyses in lamprey typically focus on the larger Muller reticulospinal axons as these allow stable intracellular recordings to be made from identified axons (Oliphint et al., 2010; Wood and Cohen 1981; McHale et al 1995; Hall et al., 1989; Zhang et al., 2011). These axons regenerate to make functional synaptic connections below lesion sites (Mackler and Selzer 1987).

While regeneration is associated with functional recovery in the lamprey, regeneration is never complete (it ranges from 0-70%; McClellan 1994) and regenerated axons project to ectopic locations (Wood and Cohen 1981): regeneration thus does not ‘repair’ the spinal cord to its original state. For regeneration to underlie locomotor recovery either requires redundancy in the unlesioned descending input to allow normal function despite the reduction in axon number, or some additional factor that can compensate for the reduced descending input. The latter seems more likely given that redundant descending inputs would be costly to develop and maintain, and given the evidence from the lamprey to human spinal cord of functional changes below spinal cord lesion sites (Grasso et al 2004; D’Amico et al 2014; Parker 2017).

Morphological analyses have shown that regenerated Muller axons make fewer synapses below the lesion site, and that these synapses contain fewer vesicles and have smaller active zones than axons from unlesioned animals (Oliphint et al., 2010). Despite this, Muller axons evoke postsynaptic inputs that match those in unlesioned animals (Cooke and Parker 2009). While this is unexpected given the morphological changes shown by Oliphint et al, there are compensations that can allow single axons to evoke the same postsynaptic input (see Parker 2017). While the large Muller axons are convenient targets for analyses, lesioning the medial column where they project does not abolish locomotion in functionally recovered animals. Lateral column axons instead seem to be more important for locomotor recovery (McClellan 1990; Chen et al 2017). This analysis has thus focused on these axons. The results suggest that regenerated synaptic inputs to motor neurons below the lesion site match those in unlesioned animals, but that the input is evoked using different release properties. However, inputs to motor neurons from axons above the lesion site are altered to become functionally stronger than those in unlesioned animals.

## Materials and methods

Juvenille adult lampreys *(Petromyzon marinus)* between 100-130mm were obtained from commercial suppliers (Acme Lamprey, Maine, USA). Animals were maintained in aquaria at room temperature. To lesion the spinal cord, animals were anaesthetized by immersion in MS-222 (300mg/ml, pH adjusted to 7.4). The spinal cord was exposed by making a dorsal incision approximately 1cm below the last gill and was then completely transected using iridectomy scissors. The incision site was repaired with Vet Bond Tissue adhesive (3M Animal Care Products). Following transection, animals were kept at 21 °C for 8-10 weeks (Cohen et al. 1999). All experiments were conducted under license of the UK Home Office (Animals Scientific Procedures Act 1986) and with the approval of the local ethical committee.

Video and electromyogram (EMG) recordings were made from control animals and animals 8-10 weeks after lesioning to assess the degree of recovery. The animals used here were assessed to have either recovered normal or near-normal locomotion (‘good’ recovery) or failed to show any recovery (‘poor’ recovery; see Hoffman and Parker 2011 for details) The spinal cord was then isolated for intracellular recordings by anaesthetising animals in MS-222 and removing the spinal cord and notochord. The spinal cord was isolated from the notochord and placed ventral side up in a Sylgard-lined chamber where it was superfused with Ringer containing (in mM): 138 NaCl, 2.1 KCl, 1.8 CaCl_2_, 1.2 MgCl_2_, 4 glucose, 2 HEPES, 0.5 L-glutamine. The Ringer was bubbled with O_2_ and the pH adjusted to 7.4 with 1M NaOH. The experimental chamber was kept at a temperature of 10-12°C. Calcium was reduced to 50% in low calcium Ringer and increased to 200% in high calcium Ringer (Parker 2003); changes in glucose levels were made to maintain osmolarity.

Paired recordings were made from axons in the lateral tract and motor neurons using thin-walled micropipettes filled with 3M potassium acetate and 0.1M potassium chloride. The lateral tract in the lamprey spinal cord also contains axons from the giant interneurons which relay sensory input to the brainstem and reticulospinal neurons. Giant interneuron axons are located near the lateral edge of the spinal cord whereas the reticulospinal axons are in the inner two-thirds of the tracts (Rovianen 1979): recordings were thus made on the medial edge of the lateral tract to avoid giant interneuron axons. Motor neurons were identified by recording orthodromic extracellular spikes in the corresponding ventral root following current injection into their somata. Axons were identified by recording antidromic extracellular spikes on the rostral end of the spinal cord following their stimulation, and by the absence of a slow afterhyperpolarisation following the action potential. Monosynaptic connections were identified by their reliability and constant latency following presynaptic stimulation at 20Hz (Berry and Pentreath 1976). To minimise potential differences due to the location of cells in different regions of the spinal cord, all experiments were performed in the rostral trunk region (i.e. the first 2cm of the spinal cord immediately caudal to the last gill). Motor neurons in lesioned animals were recorded approximately 2-3 segments above or below the lesion site (a segment is defined by the presence of a ventral root). Axons were recorded approximately 5 segments above the lesion site. An Axoclamp 2A amplifier (Axon Instruments, California) was used for voltage recording and current injection. Where necessary, the membrane potential in control and in altered Ringer solutions was kept constant by injecting depolarising or hyperpolarising current using single electrode discontinuous current clamp. Data were acquired, stored, and analysed on computer using an analogue-to-digital interface (Digidata 1200, Axon Instruments, California) and Axon Instruments software (pClamp 8).

Axon spikes were evoked either by injecting 1ms depolarizing current pulses of 10-60nA, or on rebound from hyperpolarizing current pulses (2-5ms, 1-5nA). Single synaptic inputs were evoked ten times at 0.1Hz (no activity-dependent plasticity occurs at this frequency; Brodin et al 1994) and the input averaged to measure the properties of the connection. The plasticity of inputs during spike trains was examined using presynaptic stimulation at 20Hz for 1s (Brodin et al 1994): a single input was evoked at 500ms, 1s, 2s, and 3s after the end of the train to monitor the recovery from any plasticity. Ten spike trains evoked at 30 s intervals were averaged to determine the properties of the connection. The initial EPSPs in the trains were also used as a measure of low frequency-evoked inputs.

EPSP amplitudes were measured as the peak amplitude above the baseline immediately preceding the spike. At 20Hz there was little summation of EPSPs during spike trains and thus inputs could be measured without correction (Parker 2003). The initial EPSP, the paired pulse (PP) plasticity, and the plasticity over the 2^nd^ to 5^th^ spikes (Train_2-5_), the 6^th^ to 10^th^ spikes (Train_6-10_), and the 11^th^ to 20^th^ spikes in the train (Train_11-20_) were measured. Depression was defined as a reduction to at least 90% and facilitation to at least 110% of the initial EPSP amplitude: when depression or facilitation did not at least reach these levels the connection was classified as unchanged. Paired pulse plasticity was expressed as EPSP_2_/EPSP_1_, and plasticity over different regions of the spike train as EPSP_Train_/EPSP_1_. Single, low frequency-evoked EPSPs were used to measure EPSP rise times and half-widths.

Values are presented as mean and SEM. Statistical significance was examined using two-tailed paired or independent t-tests, or one-way analysis of variance (ANOVA), and differences in the proportion of effects by Chi square. When an ANOVA was used a Tukey test was used for post hoc analysis of differences between groups. N numbers in the text refer to the number of connections examined. No more than three connections were examined in a single spinal cord. Statistical tests were only performed when the sample size allowed a power of at least 0.8: where a p value is not reported the sample size was below that needed for this power.

## Results

### Basic synaptic properties

In unlesioned animals, the initial EPSP amplitude was 1.05 ±0.09mV and the paired pulse ratio 1.1 ±0.05 (n=62; Figure 1A,B). In lesioned animals, the EPSP amplitude, rise time, and half-width above (n=19) and below (n=15) the lesion site did not differ significantly to unlesioned values (p>0.05; Fig. 1A,B,C). However, the paired pulse ratio was significantly higher above the lesion site (1.42±0.14) than for EPSPs in unlesioned animals or below the lesion site (1.1±0.05 and 1.1±0.09, respectively; p<0.05; Fig. 1B), an effect that can be associated with a lower release probability (Zucker and Regehr 2002). EPSPs can also show an electrical component or be exclusively electrical. Morphological analyses of medial column Muller axons suggest a reduction of electrical connections after injury (Wood and Cohen 1981), but the proportion of electrical vs non electrical synapses here did not differ significantly between unlesioned and lesioned animals above and below the lesion site (data not shown; p>0.05, Chi square). There were no changes in axonal action potential properties that could influence transmitter release (e.g. spike broadening; Parker et al 1997; data not shown). These changes have been reported in regenerated axons in lamprey, but they have returned to unlesioned values at the time the analyses here were performed (McClellan et al 2008).

**Figure 1:**
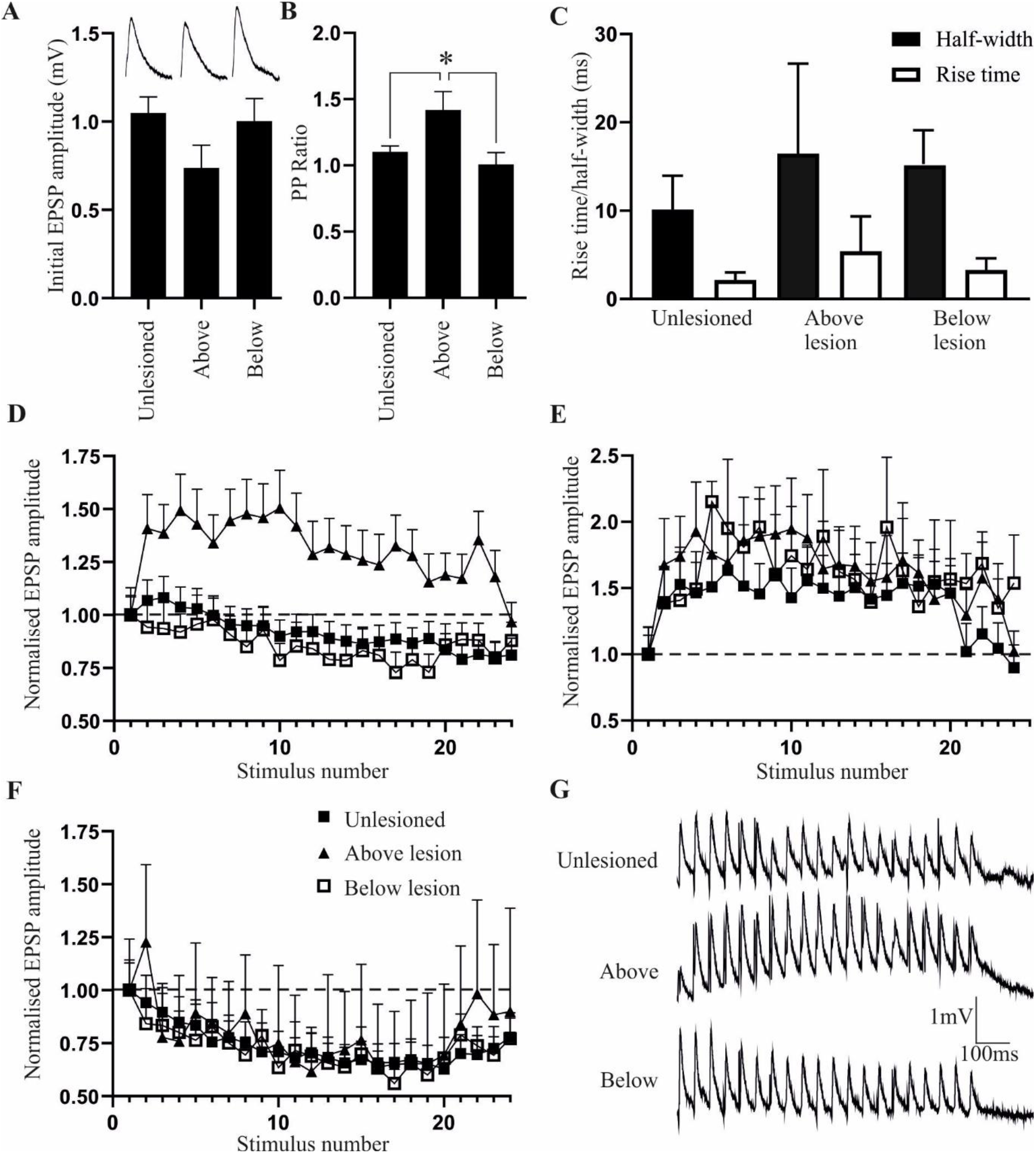
Basic synaptic properties in unlesioned and lesioned spinal cords above and below the lesion site. The amplitude (A), paired pulse (PP) ratio (B) and half-width (C) of single low frequency-evoked EPSPs. The inset in (A) shows averaged (n=10) low frequency-evoked EPSPs in the different conditions. (D) The averaged activity-dependent plasticity of all synaptic inputs in unlesioned, above, and below lesion cords over 20Hz spike trains, and the recovery of the effect after the end of stimulation (stimulus numbers 21-24). (E) Comparison of connections that only showed facilitation over spike trains. (F) Comparison of connections that only showed depression over spike trains. (G) Traces showing averaged (n=10) inputs over spike trains in an unlesioned spinal cord, and above and below the lesion site.

### Activity-dependent plasticity

Facilitation was the usual paired pulse effect (i.e. the 2^nd^ EPSP compared to the 1^st^) to 20Hz stimulation in both lesioned and unlesioned animals. Plasticity over spike trains was assessed from the effect over the 11^th^ and 20^th^ EPSPs in response to 20Hz stimulation (effects typically plateau over this part of the spike train; Fig. 1D). In unlesioned axons, the average effect was depression to approximately 80% of the initial EPSP amplitude. However, individual connections varied (see also Brodin et al 1994). In unlesioned animals, 30 (of 62) connections depressed, 14 connections facilitated, 2 connections were biphasic, and 16 were unchanged (i.e. showed no activity-dependent plasticity over the spike train). In lesioned animals, below the lesion site depression was again the usual effect (10 connections depressed, 2 facilitated, and 3 were unchanged). In contrast, above the lesion site facilitation was the most common effect over spike trains (Fig. 1D; 12 connections facilitated, 4 connections depressed, and 3 were unchanged). While the proportions of the different types of activity-dependent plasticity did not differ in unlesioned and below lesion spinal cords, above the lesion site facilitating connections were significantly more common (p<0.05, Chi square).

When connections were separated by the type of activity-dependent plasticity, the degree of facilitation did not differ in unlesioned and above and below lesion spinal cords: facilitation developed to approximately 150% over Train_11-20_ in all cases (Fig. 1E,G), and did not differ significantly in unlesioned and above lesion axons (p>0.05; the number of facilitating connections below the lesion site was too small to compare statistically). Depression also did not differ in unlesioned and above and below lesion spinal cords (Fig. 1F,G), the magnitude not differing significantly in unlesioned and below lesion axons (p>0.05; the number of depressing connections above the lesion site was too small to compare statistically).

In unlesioned animals there was no significant relationship between the initial EPSP amplitude and the paired pulse (PP) ratio (r^2^=0.02) or plasticity over the spike train (Train_2-5_ plasticity (r^2^=0.03), Train_11-20_ plasticity (r^2^=0.08); n=62; Figure 2A). The same applied to the above lesion PP ratio (r^2^=0.02) and Train_2-5_ (r^2^=0.03), and Train_11-20_ plasticity (r^2^=0.07; Figure 2B). The initial EPSP amplitude thus did not predict the type of activity-dependent plasticity at these connections (a similar effect occurs at other excitatory synapses in the lamprey spinal cord; Parker 2003a). However, below the lesion site, while there was no significant relationship between the initial EPSP amplitude and the paired pulse ratio (r^2^ =0.16), there were significant negative relationships with the Train_2-5_ (r^2^=0.23; data not shown) and Train_11-20_ plasticity (r^2^=0.29; Figure 2C). Larger initial EPSPs thus evoked greater depression, an effect that can be associated with release probability-dependent depression due to depletion of greater postsynaptic desensitisation (Zucker and Regehr 2002).

**Figure 2:**
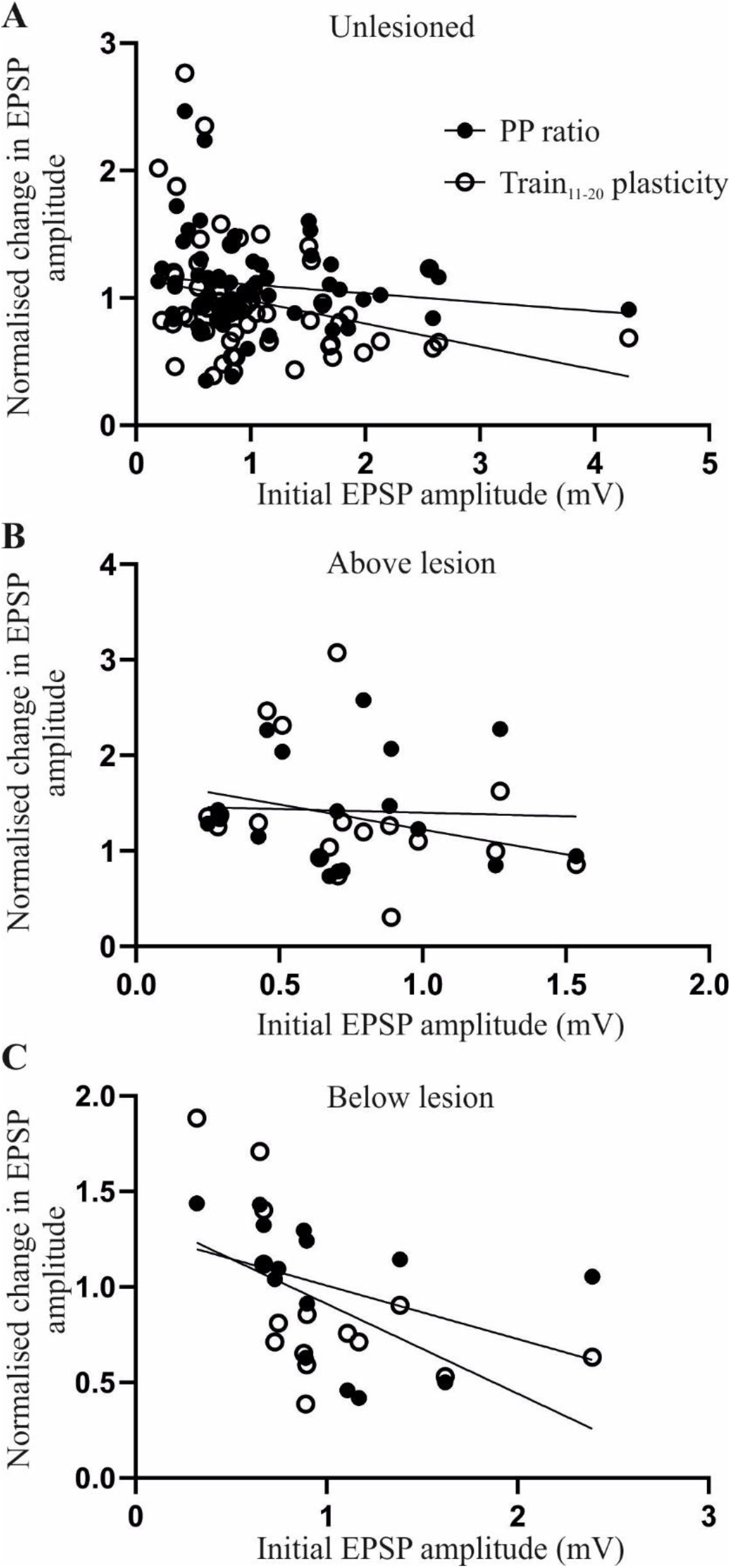
The relationship between the PP or Train_11-20_ plasticity and the initial EPSP amplitude in unlesioned spinal cords (A), above the lesion site (B), and below the lesion site (C). All graphs show linear regression fits of the EPSP amplitude to the two regions of plasticity over the spike train. Only the PP ratio and Train_11-20_ plasticity is shown for clarity.

### Release properties

Synaptic properties reflect the parameters of transmitter release, including the number of vesicles or vesicle release sites, the release probability, and the postsynaptic response to transmitter (i.e. quantal amplitude; Zucker and Regehr 2002). The number of vesicles was estimated from the model of Wang and Zucker (1998):

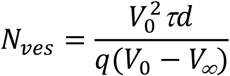

*V_o_* is the initial EPSP amplitude, *τd* the inverse rate constant of EPSP decay (expressed as the number of presynaptic spikes needed for the EPSP to fall to *1/e* of the initial value), *q* the mean quantal amplitude (assumed to be 0.1mV; Parker 2003), and *V_∞_* the EPSP amplitude at the plateau level of depression. Because this analysis depends on the rate of depression (Wang and Zucker 1998; Schneggenburger et al 1999; Millar et al 2002), it could only be applied to unlesioned and below lesion axons as the number of depressing connections was too small above the lesion site. The analysis also required that the input decreased to *1/e* of the initial value over the spike train. This often does not happen even over longer spike trains (>50 EPSPs), probably due to the concomitant development of activity-dependent replenishment of the releasable pool over the spike train (Parker 2000). As a result, the extrapolated exponential depression calculated from the initial depressing EPSPs in the train was used to determine *τd* and *V_∞_* to allow approximation of the vesicle pool (see Parker 2003). Any influence of replenishment would reduce the rate of rundown and give an overestimation of the initial vesicle pool. This would apply equally to unlesioned and lesioned synapses unless there was a difference in the rate of replenishment between unlesioned and lesioned synapses (Parker 2000), an effect that would be functionally equivalent to an increase or decrease in vesicle numbers if the rate of replenishment was faster or slower, respectively. In unlesioned animals the mean number of vesicles was 490±90, which is within the range of direct vesicle counts of 100-500 vesicles in the middle section of the vesicle pool from Muller axons in lamprey (Gustafsson et al 2002). However, below the lesion site the estimated mean number of vesicles was almost 50% greater (717±100; p<0.05; Fig. 3A,B).

**Figure 3:**
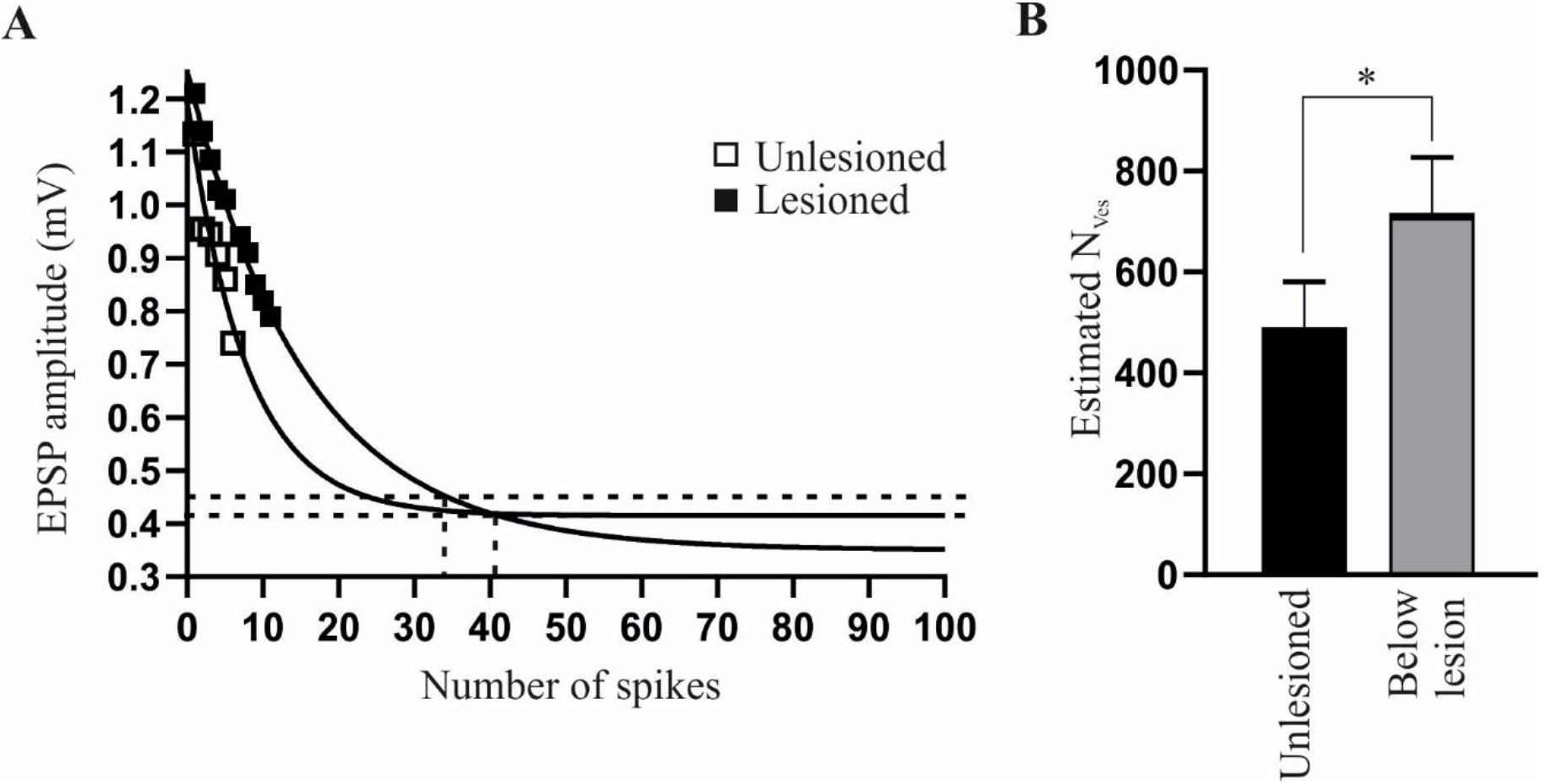
Estimate of the number of vesicles (N_Ves_) at connections in unlesioned spinal cords and connections below the lesion site. (A) Graph showing the extrapolated exponential decay of the EPSP in the two conditions, used to derive parameters to estimate N_Ves_ (see text for details). (B) Graph showing the N_Ves_ in unlesioned and below lesion connections.

Presynaptic or postsynaptic parameters were assessed using the coefficient of variation (CV^-2^) to compare changes in mean and variance of the initial and 20^th^ EPSPs in spike trains (Brock et al 2020). The analysis assumes that parallel changes in synaptic amplitude and CV^-2^ reflect presynaptic effects, while synaptic changes without a change in CV^-2^ reflect postsynaptic effects (Faber and Korn 1991). As the method is sensitive to noise (Brock et al 2020), only connections with no spontaneous synaptic activity were analysed. In unlesioned animals, 3of 6 facilitating connections had CV^-2^ changes associated with presynaptic effects, 3 with postsynaptic or mixed pre and postsynaptic effects: for depressing connections 17 had CV^-2^ changes consistent with a presynaptic effect and 11 with postsynaptic/mixed effects. Above the lesion site, 4 of 8 facilitating connections had changes consistent with a presynaptic effect, 4 with mixed or postsynaptic effects. Below the lesion site 4 connections showed CV^-2^ changes consistent with presynaptic effects and 2 with mixed or postsynaptic effects. The proportions of presynaptic and postsynaptic/mixed effects did not differ in unlesioned or lesioned spinal cord (p>0.05 Chi square).

As the CV^-2^ method relies on certain conditions and assumptions to identify a pre or postsynaptic locus of change (Faber and Korn 1991; Brock et al 2020), release properties were also examined using the variance-mean method (Clements and Silver 2000). This examines postsynaptic potentials in normal, high, and low calcium Ringer. Unless the release probability is low (<0.3), the relationship of the EPSP variance and mean in different calcium Ringers results in a parabolic relationship from which release parameters can be estimated (Clements and Silver 2000 for details): *N_min_*, the number of release sites is determined from the width of the parabola; *p*, the probability of release from the curvature; and *q*, the postsynaptic quantal amplitude from the initial slope (Fig. 5Ai).

**Figure 4:**
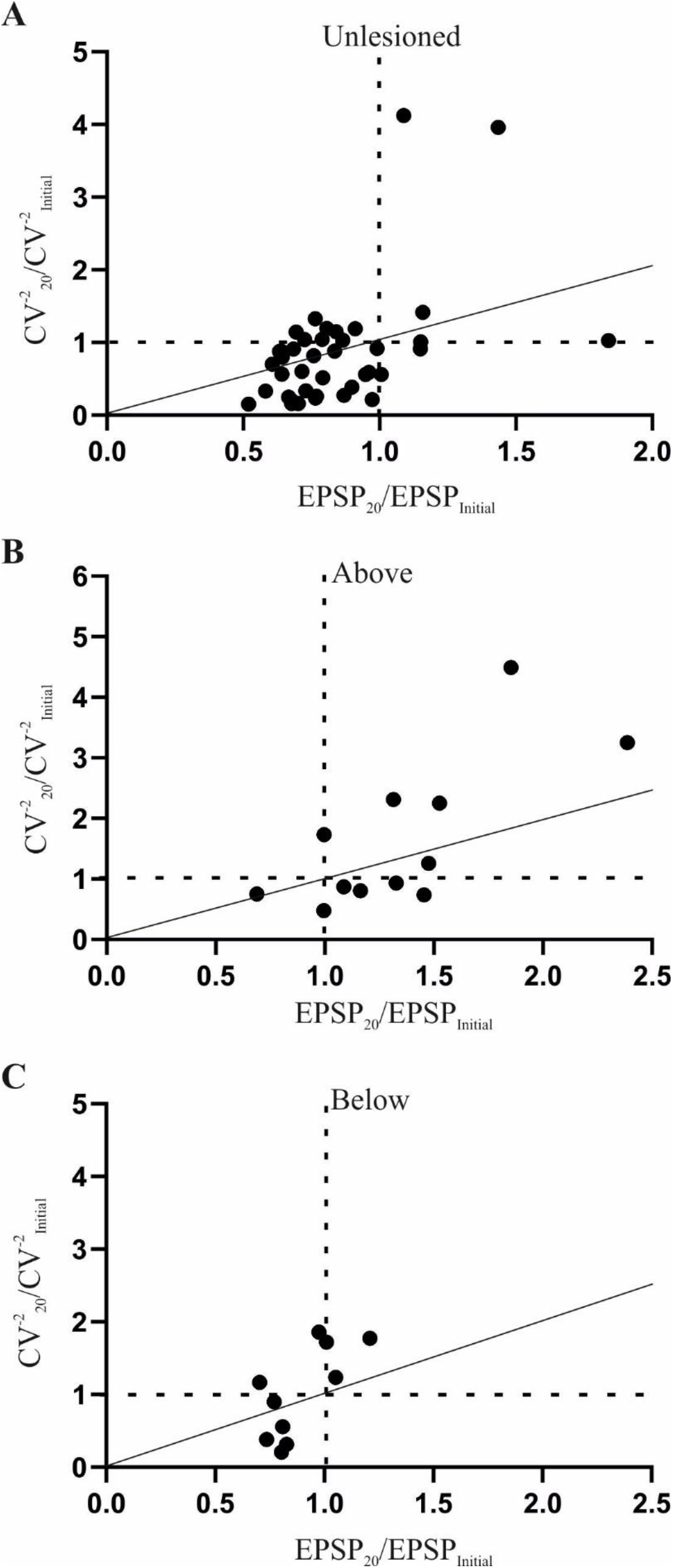
The analysis of the plasticity over the spike train ((EPSP_20_/EPSP_1_) and the inverse of the coefficient of variation (CV^-2^_20_/CV^-2^_1_) for unlesioned (A), above lesion (B), and below lesion connections (C). Note that the *x* axis shows the plasticity over the spike train and the *y* axis the change in the coefficient of variation. A presynaptic change in plasticity is indicated by values falling either on or below (for depression) or above (for facilitation) the diagonal line, a postsynaptic change when values fall on the horizontal dashed line (i.e. no change in the CV^-2^_20_/CV^-2^_1_ despite a change in the EPSP), and both pre and postsynaptic changes for depression and facilitation when values fall above and below the diagonal line, respectively (see Faber and Korn 1991).

**Figure 5:**
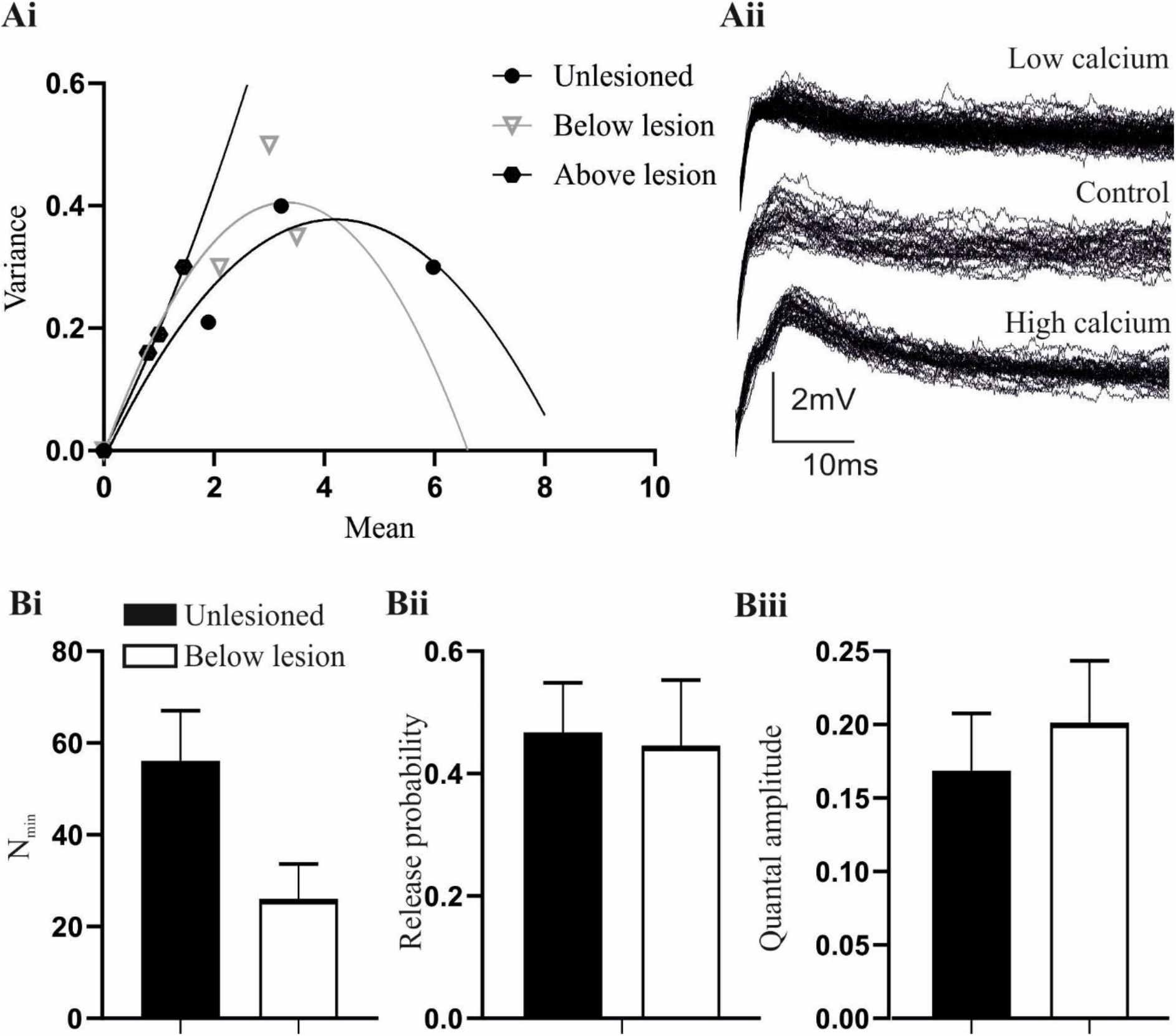
(Ai) A variance-mean analysis of connections in unlesioned spinal cords, and above and below the lesion site (see text for details). (Aii) Sample traces in an unlesioned spinal cord in normal and low and high calcium Ringer. (Bi) The estimated number of release sites (N_min_) of unlesioned and below lesion connections. (Bii) The estimated release probability of unlesioned and below lesion connections. (Biii) The estimated quantal amplitude of unlesioned and below lesion connections.

The analysis again only includes connections that were stable throughout the experiment and in which EPSP amplitudes could be measured unequivocally (i.e. without contamination by an electrical component to the EPSP or spontaneous synaptic inputs). This was a significant issue, as the lateral tract axons are relatively small and recordings stable enough to allow the effects of the different calcium Ringer solutions to be examined were not common, and low calcium Ringer often increased background synaptic inputs, presumably through a reduction of surface screening (Piccolino and Pignatelli 1996). This meant that the *n* number of fully analysed connections is small (this is not uncommon for variance-mean analyses; see for example, Kazama and Wilson 2008; Lawrence et al 2015; Mitchell and Silver 2000; Malagon et al 2016). In regenerated axons that evoked depression below the lesion site *N_min_* was generally reduced (n=3; Fig. 5Bi), consistent with the sparser anatomical connectivity reported for medial column axons (Oliphint et al 2010), but *p* was unchanged compared to synapses in unlesioned cords (n=3; Fig. 5Bii), an effect consistent with the similar activity-dependent depression over spike trains (see Fig. 1F,G). However, *q* was increased (n=3; Fig. 5Biii), suggesting a potential change in postsynaptic response to transmitter. Analyses of facilitating connections above the lesion site consistently showed a linear rather than parabolic relationship between the variance and mean (n=4; Fig. 5Ai). This is indicative of a low release probability (<0.3; Clements and Silver 2000), and is consistent with the higher PP ratio and the development of facilitation above the lesion site (Fig. 1B,D). The absence of the parabolic relationship prevented estimation of *N_min_* and *p*, but the slope of the relationship was not increased compared to unlesioned animals, suggesting against a change in *q* above the lesion site (Clements and Silver 2000).

### Influence of degree of recovery

Poorly recovered lampreys show an altered relationship between regenerated inputs and spinal cord excitability compared to animals that recover well (Hoffman and Parker 2011), suggesting that the interaction of regenerated synapses and their spinal cord targets, rather than just the presence of regeneration, is important to recovery. An obvious consideration in determining the functional relevance of the synaptic properties examined here is to examine the properties of regenerated synapses in animals that recovered poorly. However, while this has been done for other aspects after lesioning in lamprey (Hoffman and Parker 2011; Becker and Parker 2015), this was difficult in this analysis as poor recovery tends to be associated with the absence of regeneration, which limited the sample size of paired recordings. However, synaptic properties were examined in animals that recovered poorly despite the presence of regeneration. The number of these animals, and the sample size of connections, is low and thus the data is preliminary. Connections below the lesion site had smaller amplitudes than connections in animals that showed good recovery (0.23±0.15mV (n=4), respectively; Fig. 6A), usually depressed to a greater extent, and uncharacteristically for these synapses tended to exhibit failures. Above the lesion site (n=7), connections again showed facilitation albeit from smaller initial EPSP amplitudes than connections that recovered well (0.57mV±0.06mV; Fig. 6A,B).

**Figure 6:**
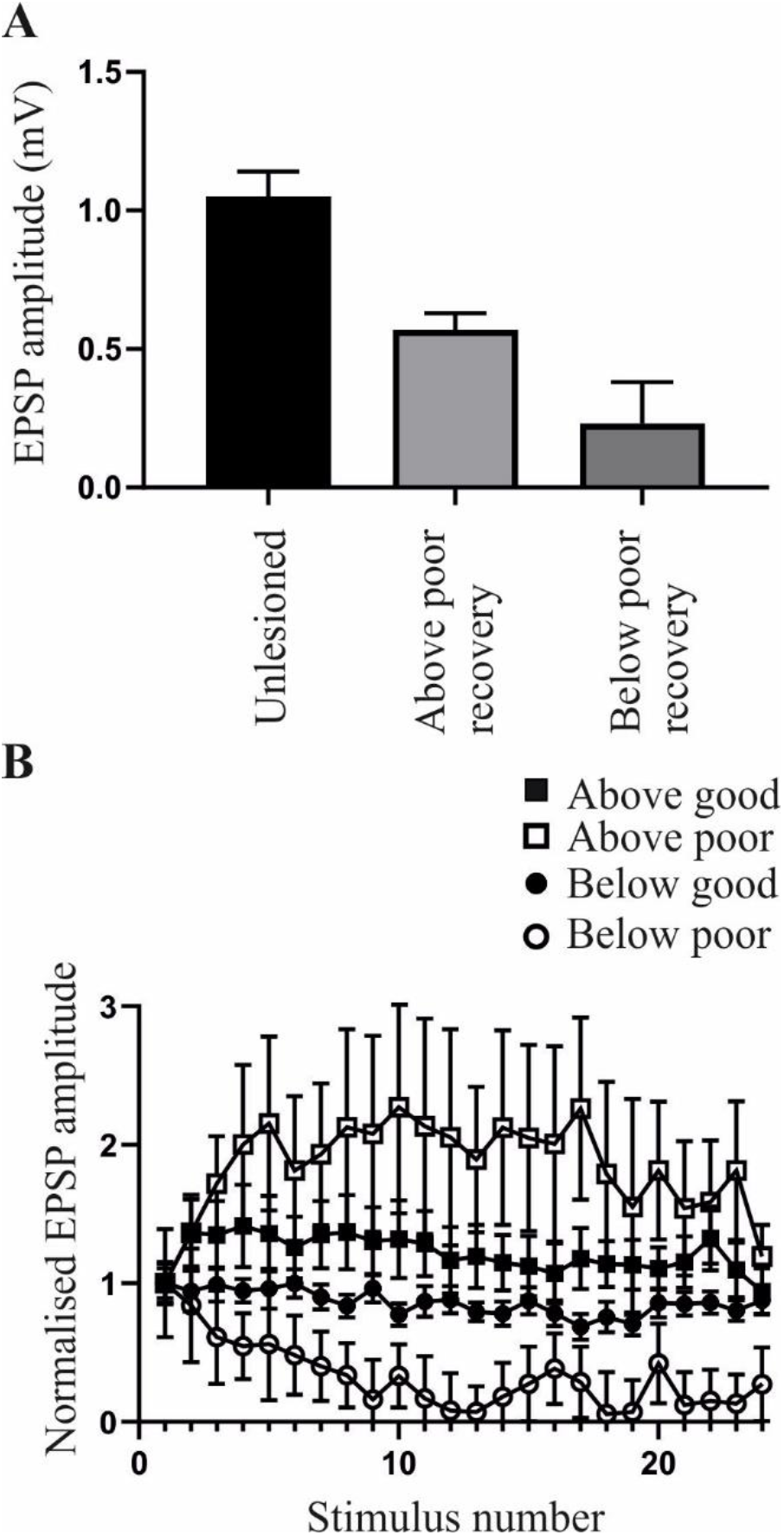
Differences in synaptic properties in animals that recovered well or poorly. (A) The EPSP amplitude below the lesion site in good and poor recovery. (B) The activity-dependent plasticity of connections above and below the lesion site in animals that recovered well or poorly.

## Discussion

This analysis has examined the properties of regenerated axons in the lamprey. Synaptic inputs from lateral column reticulospinal axons below the lesion site matched those from unlesioned axons in terms of their amplitude and activity-dependent plasticity. However, despite the same output being generated, release properties differed. In contrast, connections made by axons above the lesion site differed to those in the unlesioned spinal cord as they showed facilitation that developed from the same initial EPSP amplitude and would thus be functionally stronger.

Regeneration remains the dominant approach in spinal cord injury research. Analyses focus on the anatomical properties of regenerated axons (e.g. their number, location, and how far they regenerate below the lesion site), and the correlation of regeneration with recovery. However, it is not enough just for axons to regenerate: functional recovery requires that they make appropriate connections with targets below the lesion site. Differences in the release properties that determine the features of these connections could thus contribute to the variable influence of regeneration on recovery seen in experimental model systems like the lamprey (Parker 2017) and in clinical trials of regeneration (Steward et al 2012).

It is difficult to examine synaptic properties directly to compare effects in unlesioned and lesioned spinal cords. A range of approaches were used here, including direct measurements of basic synaptic properties, the estimation of vesicle numbers, comparison of presynaptic vs postsynaptic mechanisms (1/CV^2^), and comparison of release properties (variance-mean analysis). While these are commonly used approaches, they have limitations and determining the mechanisms of release at central synapses is still difficult (Lanore and Silver 2016; Pulido and Marty 2017). Changes in postsynaptic response can be assessed simply and directly by comparing the response to exogenously applied glutamate in functionally isolated cells (e.g. in the presence of TTX; Parker et al 1997), but this only works for comparisons of the same cell under different conditions (e.g. before and after application of a neuromodulator), not for comparing effects in different spinal cords. The variance-mean approach makes fewer assumptions than traditional quantal mechanisms (Clements and Silver 2000), including the CV^-2^ analysis (Faber and Korn 1991), and has been applied at a range of central synapses (see Lanore and Silver 2016). However, the analysis needs relatively long-term stable recordings from relatively small presynaptic axons to allow the changes in Ringer calcium levels, and evoked EPSPs that are not contaminated by spontaneous background inputs, requirements that limit the sample size. However, the results of the various analyses tended to be consistent with each other, providing support for the general conclusions.

The same amplitude and activity-dependent properties of regenerated axon inputs to motor neurons below the lesion site as unlesioned animals suggests against an increase in the synaptic strength of individual axons to compensate for the overall reduction of the descending input after recovery from a spinal cord lesion (fully recovered animals show at least a 30% reduction in the number of axons; Cohen et al 1988; McClellan 1990). However, the same synaptic effect is evoked by different synaptic mechanisms. The different relationship between the initial EPSP amplitude and the paired pulse ratio in below lesion synapses suggests a change in presynaptic release (that larger initial EPSPs evoked greater depression may reflect a larger quantal content with a concomitant increase in vesicle depletion and depression or greater postsynaptic desensitisation; Zucker and Regehr 2002), while the variance-mean analysis suggests an increase in the postsynaptic response (quantal amplitude, *q*) below the lesion site, an effect that is consistent with previously identified sub-lesion functional changes (Parker 2017). This effect could scale up regenerated inputs to allow them to match the properties of unlesioned synapses despite the reduced number of release sites suggested by the reduced *N_min_* from the variance-mean analysis, and the potential for a reduced number of synaptic contacts from morphological analyses of regenerated Muller axons below the lesion site (Oliphint et al 2010). The increase in the estimated vesicle numbers below the lesion site could also help to strengthen each contact. The combination of these presynaptic and postsynaptic mechanisms below the lesion site may account for the lack of a difference in the proportions of putative presynaptic and postsynaptic in the CV^-2^ analysis.

Why should different release properties be used to generate the same synaptic output below the lesion site? The maintenance of unlesioned properties could obviously reflect the need for inputs of a certain magnitude to effectively activate sub-lesion networks. The difficulties of recapitulating the original development of these axons and their synaptic contacts in a mature nervous system may account for the smaller and limited number of synaptic contacts below the lesion site (Oliphint et al 2010). Each contact made by a single axon could be made stronger by increasing the transmitter release probability, but unless transmitter replenishment was also more efficient this could lead to a release probability-dependent increase in depression that would weaken inputs over spike trains (Zucker and Regehr 2002). Instead of strengthening the connection, an increase in release probability alone would thus redistribute the input over the spike train, from being initially stronger to being subsequently weaker than unlesioned inputs, depending on how many inputs were evoked. Maintaining high levels of release and the associated need for increased transmitter clearance and replenishment would also invoke a significant energy cost (Howarth et al 2012), which may be an issue for axons that are still regenerating and given the compromised state of the lesioned spinal cord. The scaling of the synapse through a postsynaptic increase in quantal amplitude and presynaptic increase in vesicle numbers suggested here could maintain the pre-lesion synaptic amplitude at the unlesioned value over spike trains despite each axon potentially making fewer synaptic contacts, without the energetic costs of upregulating transmitter release and replenishment mechanisms. The various changes in cellular and synaptic properties seen below the lesion site in various model systems from lamprey to mammals could make the spinal cord below the lesion site more excitable (D’Amico et al 2014; Parker 2017). These postsynaptic changes may better compensate for the reduced descending input below lesion sites than a presynaptic modification of the regenerated axons, as postsynaptic cells will be able to modify their response to synaptic inputs depending on the integrated descending input that they receive (Parker and Grillner 2000).

The variance-mean analysis and paired pulse ratio suggest that the release probability is reduced at synapses between lateral tract axons and motor neurons above the lesion site; this is consistent with the increased PP ratio and facilitation over spike trains in this region. However, the reduction of release probability was not associated with a significant reduction of the initial EPSP amplitude in the spike trains, which would be expected of a reduction in release probability alone, suggesting that other factors are also altered. This could reflect a concomitant increase in *q* (see Bevan and Parker 2004 for an example), but the variance-mean and CV^-2^ analyses suggests against this. It could alternatively reflect an increase in vesicle numbers (see Bevan and Parker 2004): this will need anatomical analyses as the method used here for estimating vesicle numbers is based on the rate of depression and thus cannot be applied to facilitating connections. Irrespective of the mechanism, facilitation without a significant reduction of the initial EPSP amplitude would make the summed input above the lesion site functionally stronger than connections in the unlesioned spinal cord or below the lesion site, a compensation that may allow stronger propriospinal activity across the lesion site (Courtine et al 2008).

Regeneration is never complete in lamprey, varying between 0-70% of the unlesioned value (Cohen et al 1988; McClellan 1994). That the same functional output can be generated despite a reduction in descending input either suggests that some of the descending input is redundant, which seems unlikely given the resources that would have to be dedicated to developing and maintaining these connections, or that there is some compensation for the reduction. Each regenerated axon could have increased its postsynaptic effect in proportion to the reduction of the total descending input to maintain the same overall sub-lesion synaptic input, but functionally stronger connections occurred above, not below the lesion site where a compensatory increase would be expected. However, increasing the amplitude of regenerated inputs below the lesion site to maintain the total descending synaptic at the unlesioned value would not necessarily be an effective compensation as it would not reflect the specific functional roles of inputs to different spinal cord regions. If only 50% of the axons regenerated, doubling the amplitude of each would lead to the same summed descending input, but doubling inputs to some regions would not necessarily compensate for zero inputs to other regions. Functional efficacy is not only dependent on the properties of input synaptic connections but also on the properties of the neurons and circuits that they connect to (Ullström et al 1999). Understanding the functional consequences of regeneration will thus require a focus on how regenerated axons can be made to interact appropriately above and below the lesion site.

## Notes

Conflict of interest: None

### Competing Interest Statement

The authors have declared no competing interest.

